# Effects of early life stress on biochemical indicators of the dopaminergic system: a 3 level meta-analysis of rodent studies

**DOI:** 10.1101/372441

**Authors:** V. Bonapersona, M. Joëls, R. A. Sarabdjitsingh

## 1 Introduction

During the perinatal period, the brain matures and is rapidly wired(Semple et al. 2013), rendering it particularly vulnerable to negative life experiences that might lastingly impact brain function and behavior(Levine 2005). This may contribute to the well-established observation that exposure to adverse conditions during childhood is a major risk factor for the later development of psychopathologies(Teicher et al. 2016), including schizophrenia and substance abuse(Agid et al. 1999; Enoch 2011; Dube et al. 2003; Scheller-Gilkey et al. 2002).

Prevailing evidence highlights that the dopamine (DA) system may be a prime candidate in mediating the influence of adverse events early in life on vulnerability to psychopathology(Gatzke-Kopp 2011). The DA system develops early during the embryonic period, matures throughout adolescence, and forms stable patterns during young adulthood(Money and Stanwood 2013). This prolonged development provides an extensive window of time in which adverse conditions early in life can tip the balance towards dysfunction(Money and Stanwood 2013). Indeed, alterations in this system have been consistently associated with mental disorders (for a review: (Money and Stanwood 2013)). For example, genetic variations of the DA degradation enzyme COMT are associated with schizophrenia and bipolar disorder as well as an increased risk for psychosis, autism and anxiety(Money and Stanwood 2013). In line, the DA receptor 2 is a major target for antipsychotics.

Overall, the associative studies in humans have led to the assumption that childhood adversities result in developmental alterations of the dopaminergic system. To investigate causality, preclinical studies using animal models have adopted behavioral early life stress (ELS) paradigms to mimic negative childhood conditions, aiming to understand the neurobiological substrate by which ELS adds to the development of DA system dysfunction. Although extensive, the existing literature is quite heterogeneous: it uses disparate models and outcome measures, and each study focuses on only a limited number of variables; moreover, preclinical studies are frequently underpowered(Button et al. 2013). The resulting findings are rather incoherent and difficult to interpret. This limitation hinders our understanding of the entire biological system and its development, and delays translational applicability.

To overcome these limitations, we performed a meta-analysis, a powerful method still sparsely applied to preclinical research allowing to systematically synthesize the scientific knowledge of a specific topic. Recent advances in the field of statistics such as the 3-level approach (Cheung 2014b; Assink and Wibbelink 2016) along with their implementation in R packages(Viechtbauer 2010; Cheung 2014a) now enable researchers to use more sophisticated and robust methodology when analyzing meta-data. This method allows to include multiple data-points from a single study (nesting), without necessarily knowing their (often unreported) covariance. Ultimately, this substantially increases the flexibility of meta-analysis applications and improves the validity of the conclusions drawn.

Here, we aimed to investigate whether preclinical studies support an effect of ELS on dopaminergic signaling. We included diverse types and timings of ELS models(Fig 1), and we operationalized the dopaminergic system by quantifying several biochemical markers in mice and rats(Fig 2), across brain areas(Fig 3), considering possible confounders.

**Fig 1.**
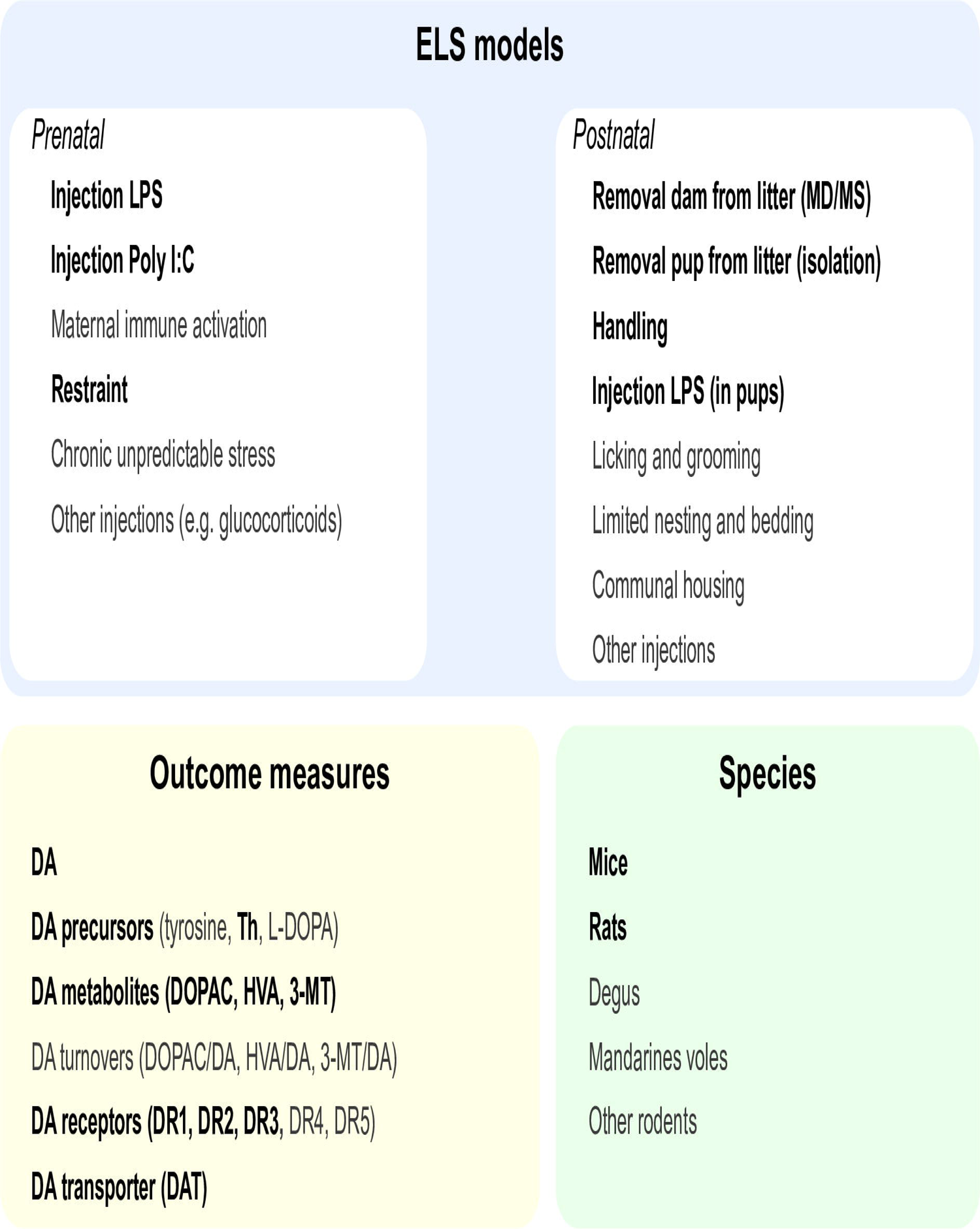
Search string and inclusion criteria. Graphical representation of the three main components of the search string. Items highlighted in bold were ultimately included in the analysis; other items were not included in the final analysis as explained in the methods’ section.

**Fig 2.**
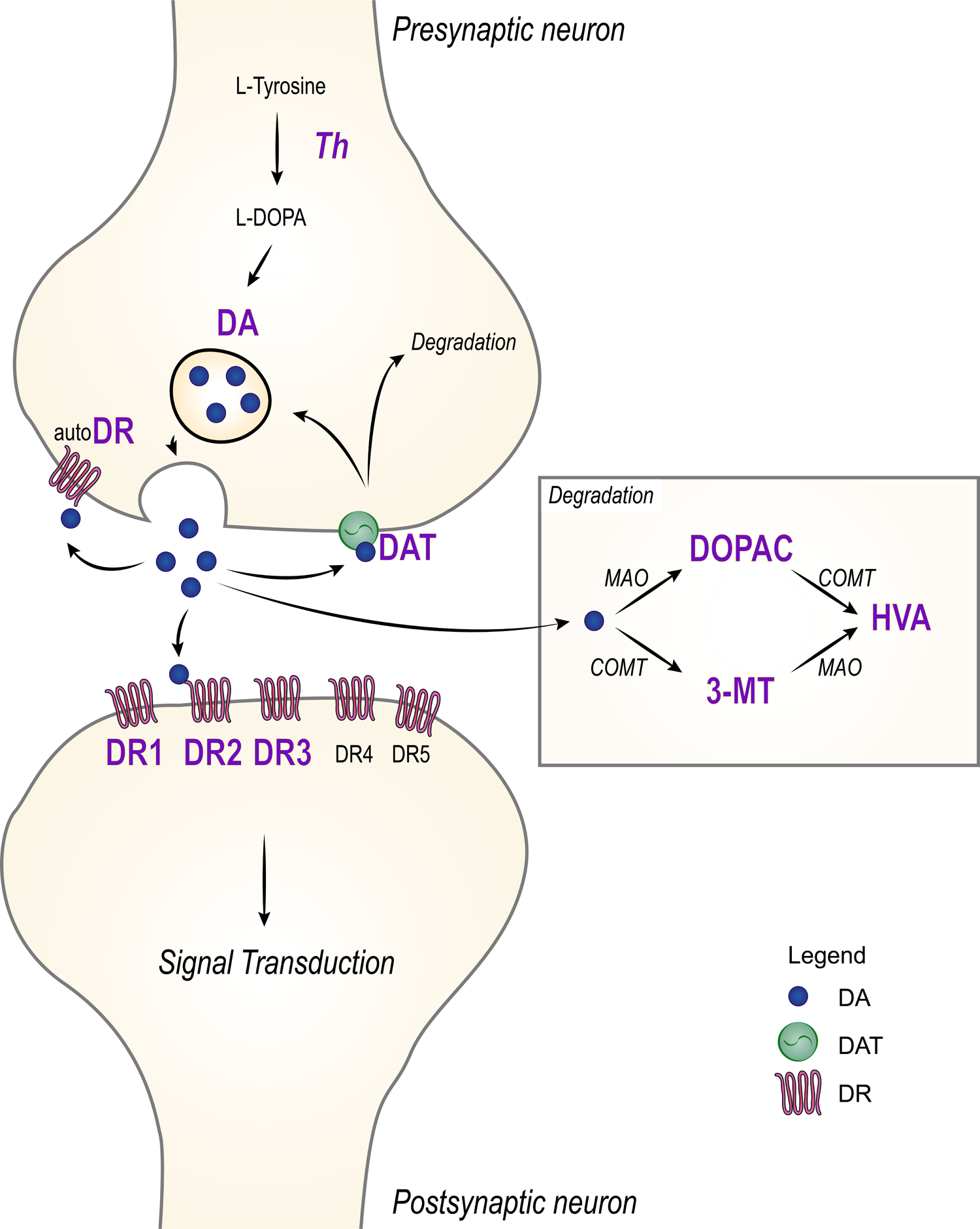
Graphical representation of signaling pathway. Dopamine is synthesized by the enzyme tyrosine hydroxylase (Th). When released in the synaptic cleft, DA can 1) bind post-synaptic receptors (DR1-DR5), 2) bind auto-receptors, 3) bind dopamine transporters (DAT), 4) be converted to the metabolites DOPAC, 3MT and HVA by the action of the enzymes MAO and COMT(Meiser, Weindl, and Hiller 2013). Items in large (purple) font were included in the meta-analysis.

**Fig 3.**
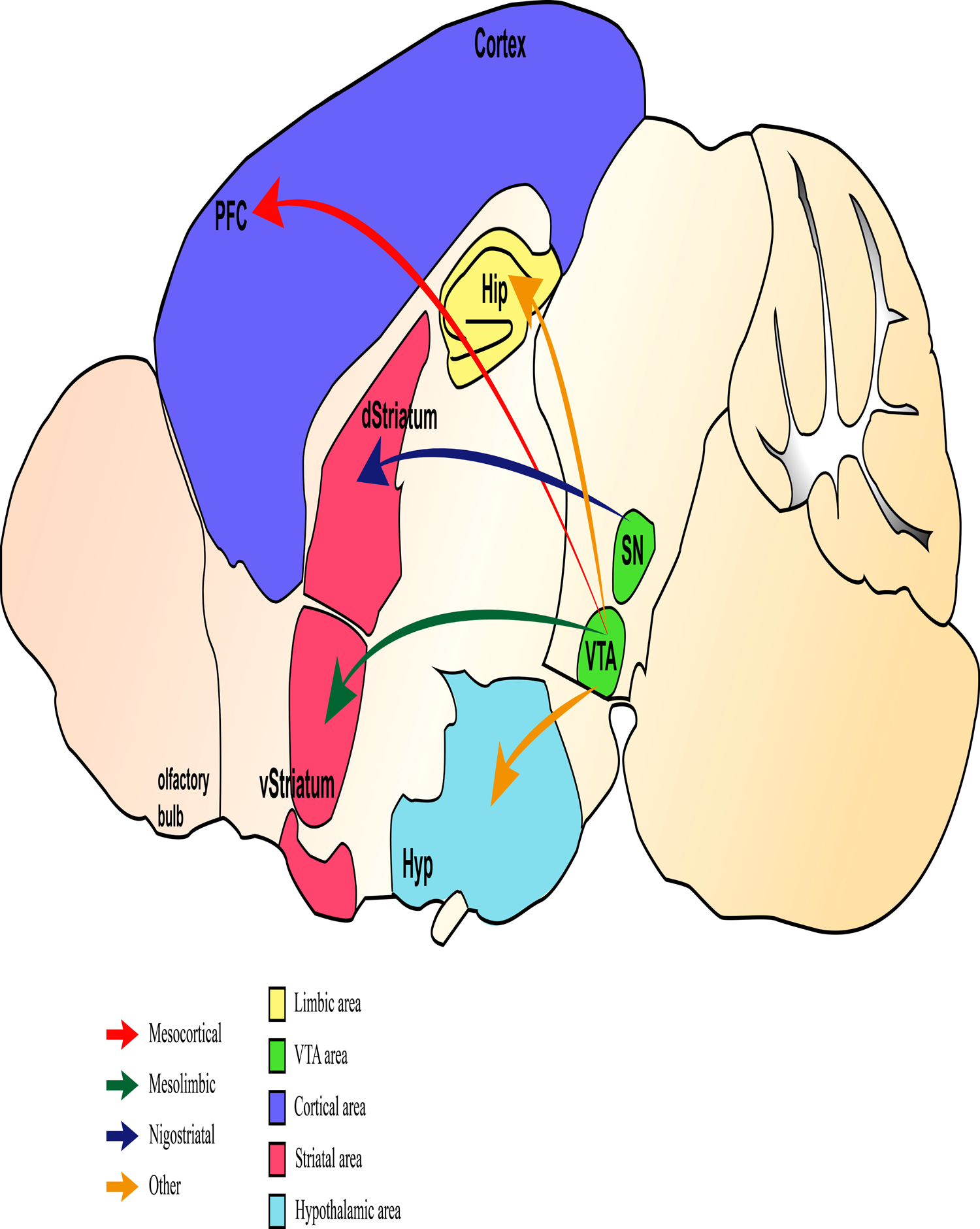
Graphical representation DA system projections. DA neurons are mainly situated in the midbrain, and can be subdivided with respect to their projection site(Grace 2016). In particular, DA neurons define separate populations of neurons that project to specific brain regions(Swanson, n.d.; Menegas et al. 2015). The major DA pathways are 1) mesocortical pathway, which defines projections from the VTA to the prefrontal cortex (PFC); 2) mesolimbic, from VTA to limbic system; 3) and nigrostriatal pathway, from substantia nigra (SN) to dorsal striatum, caudate nucleus and putamen. Other projections connect VTA to the hypothalamus, hippocampus and amygdala. Besides hosting dopaminergic neurons, these brain areas are involved in the feedback response to stress(Keller-Wood 2015).

We determined whether the quality of the studies affected the estimation of the results. To make our knowledge readily available to others, we organized all information in a freely accessible open-source dataset and created a user-friendly web-app as a tool to guide future (pre)clinical research (e.g. power analysis calculation), thereby avoiding unnecessary replication and limit animal experimentation.

## 2 Materials and Methods

The review adhered to SYRCLE (Systematic Review Center for Laboratory animal Experimentation) guidelines for protocol(De Vries et al. 2015), search strategy(Leenaars et al. 2012), and risk of bias assessment(Hooijmans et al. 2014).

### 2.1 Theoretical definitions and assumptions

We defined as “*individual comparison*” each test performed within a published study between a control group and an experimental group with a history of ELS. As often occurs in experimental studies(Aarts et al. 2014), multiple outcomes (*individual comparisons*) were collected from the same groups of animals (*nesting*).

We defined as “*experiment*” the ensemble of outcome measures from the same animals. According to this definition, each published study can report multiple experiments when conducted on different sets of animals. Similarly, experiments conducted on different sets of animals could potentially be reported in separate publications. For these reasons, we nested multiple individual comparisons belonging to the same animals within the same experiments, but considered *experiments* from the same publications as independent from each other.

### 2.2 Search Strategy

A comprehensive literature search was conducted regarding *the effects of early life stress on biochemical indicators of dopaminergic signaling* on February 14^th^ 2017. The search string was composed by the factors “dopamine”, “early life stress” and “rodents” (Fig 1 and supplementary appendix S1). The search was conducted on the electronic databases PubMed (www.pubmed.com) and Web of Science (www.webofknowledge.com). For a flow chart of the entire methodology, see Fig 4.

**Fig 4.**
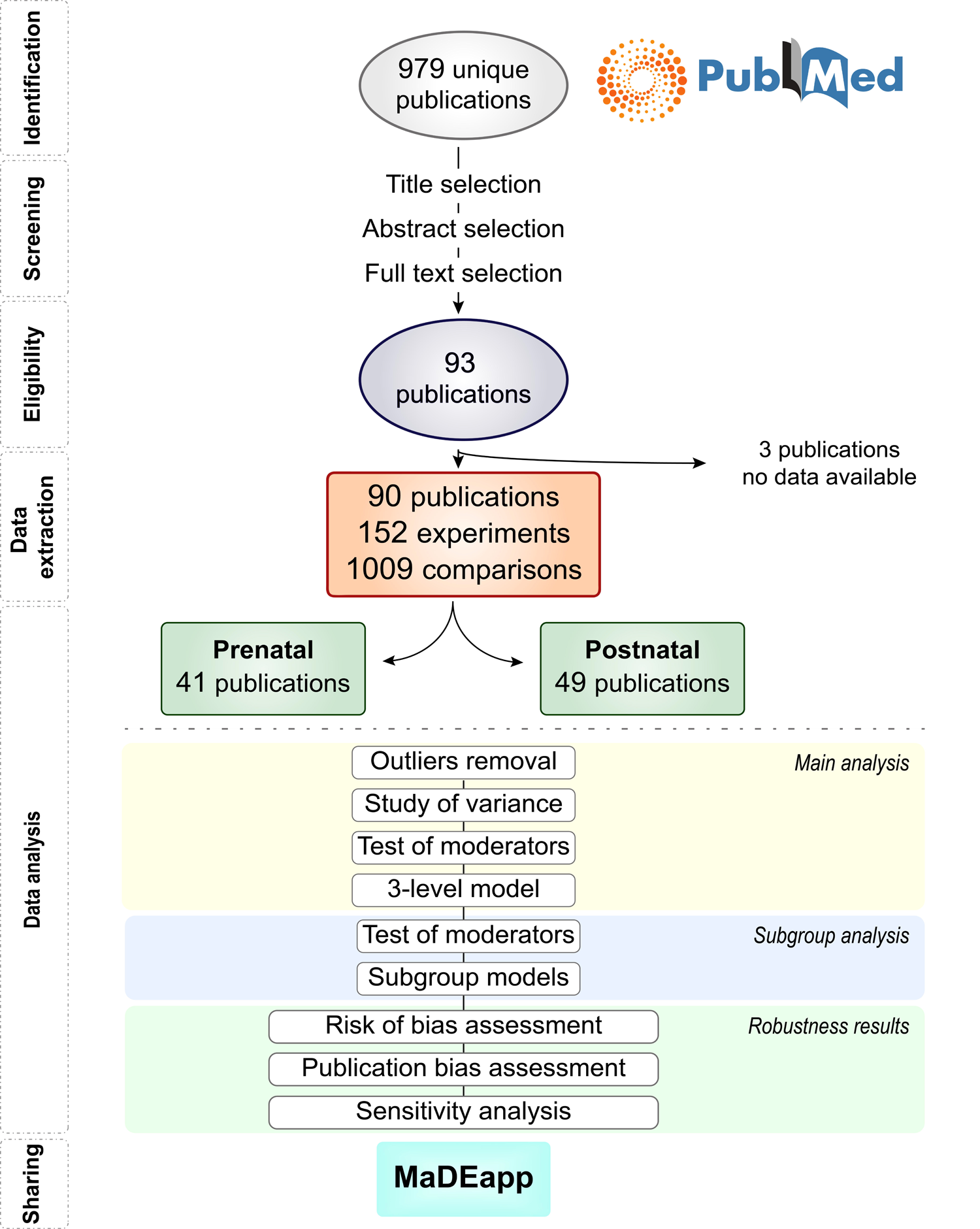
Flow-chart of study selection and analysis.

Studies’ titles and abstracts were screened, and selected if the inclusion criteria were met (supplementary Table S2-1). In case of doubt, the full text was inspected. Eligible studies were evaluated by two independent reviewers (VB and RAS).

### 2.3 Study Characteristics

#### 2.3.1 Selection and data extraction

To limit subjectivity in the data gathering and entry process, data extracted from eligible studies were recorded in a standardized database (doi:10.17632/9x9fc3v26k.1). The following information was included: species, strain, sex, type and timing (relative to age) of the ELS model, outcome, time (relative to age) of outcome, technique used for outcome, brain area investigated, number of animals used, mean, standard deviation (SD) and standard error of the mean (SEM). If only SEM was reported, SD was calculated as (Inline), where *n* = amount of animals per group. If number of animals were reported as a range (e.g. 6-8 animals per group), we used the mean of this number (e.g. 7 animals per group). If a single control group was used to compare experimental groups in which ELS was induced with different models (e.g. handling and maternal deprivation), the sample size of the control group was equally divided as control for each experimental group (e.g. n=10 in not handled control becomes n=5 for control of maternal deprivation and n=5 for control of handling)(Vesterinen et al. 2014).

When the data was not reported numerically in the publication, we contacted two authors per manuscript. If no answer was received within three weeks and after a reminder, the authors were considered not reachable. Only 5 out of 56 contacted authors replied to our request. Given the low response rate, we estimated most of the data presented only in graphs with *Ruler for Windows* (https://a-ruler-for-windows.en.softonic.com/). We tested the accuracy of this method by comparing effect sizes calculated from either supplied data or evaluated with the ruler, and verified that they were highly correlated (*R^2^* = .*74*, supplementary Figure S2-1).

Concerning metabolites, some papers reported either concentrations, turnovers or both. In 97.5% of cases it was possible to calculate concentrations from turnovers with Pythagoras. Since concentrations and turnovers are related to the same information (though not identical), only concentrations were included in the analysis in order to avoid multi-collinearity. Turnover data-points are available in the MaDEapp (*Meta-Analysis on Dopamine and Early life stress*) for consultation.

### 2.4 Assessment of risk of bias in included studies

We used the SYRCLE tool to assess the risk of bias (supplementary Table S2-2)(Hooijmans et al. 2014). The criteria are based on the possible presence of selection bias (items 1, 2 and 3), performance bias (items 4 and 5), detection bias (items 7, 8, 9) and attrition bias (item 10). Furthermore, we added the item “*quality of control*” (item 6) to the category performance bias.

Since poor reporting of experimental details plays a role in heightening the quantified risk of bias, lack of reporting was scored as *unclear* risk of bias.

For quantitative inclusion in the analysis, amount of potential bias was operationalized by summing the risk of bias of each item according to the definition: “yes” = 0, “unclear” = 1, “no” = 2. This produced a continuous variable of integer increment between 0 (no risk bias) and 20 (maximum risk of bias), which was then scaled (*mean* = 0) to interpret the studies as of average risk of bias.

### 2.5 Data synthesis and statistical analysis

#### 2.5.1 Effect size

We estimated the effect size for each individual comparison with *escalc* (R package metafor) as standardized mean difference with *Hedge’s* G method, which includes a correction factor for small sample sizes(Vesterinen et al. 2014).

#### 2.5.2 Study of heterogeneity

Heterogeneity was tested with the Cochran Q-test(Moreira et al. 2016) and I^2^ statistics(Cheung 2014b). Study of the distribution of variance was conducted for models without moderators to determine how much variance could be attributed to differences between effect sizes within experiments (level 2) and to differences between experiments (level 3). Substantial distribution of heterogeneity at these levels further encouraged the use of moderators.

#### 2.5.3 Model

We used a 3-level mixed effect model, which accounts for the anticipated heterogeneity of the studies as well as the dependency of effects within experiments. In our experimental design, the 3 levels correspond to variance of effect size between 1) animals, 2) outcomes and 3) experiments.

Since pre- and post-natal models act on different times of development that are particularly disparate regarding the array of environmental factors, we considered them as different datasets and consequently analyzed them separately.

Effect sizes were considered outliers if their z score was above +3.29 or below - 3.29(Tabachnick and Fidell 2013), and removed from the analysis.

Since we hypothesized that the effect of ELS on DA signaling may not be evident from an overall estimate, we defined a *priori* possible moderators of this effect. These belonged to two different categories: biological and technical. The biological moderators were: outcome measure used (e.g. DA and metabolites), brain area investigated, sex, species, age as a continuous variable, and whether the outcome was at a RNA level/protein level/functional (referred to as *method of assessment*). Specific regions within the brain areas were investigated only in subgroup analysis due to the limited amount of observations. We considered the type of ELS model and amount of potential bias as technical moderators. These moderators may not underlie a biological difference, but can nevertheless explain heterogeneity across studies. The postnatal ELS model ‘handling’ has been reported repeatedly to cause effects in the opposite direction to those induced by other ELS models(Levine 2005). We therefore multiplied each calculated effect size for handling by −1 (Vesterinen et al. 2014), so that the overall estimate would be in the same direction. We verified that the same conclusions would have been drawn if handling was excluded as a model (supplementary Figure S2-2).

To avoid multicollinearity among moderators, we firstly assessed each biological moderator univariately. We set the significance level at *p*<.10 to test whether a moderator significantly reduced the previously quantified heterogeneity. A less restrictive *p*-value was chosen to assure the inclusion of moderators that have a multivariate but not univariate effect(Hox 2010). Only interactions with at least 4 comparisons from 3 different publications from at least two different laboratories were quantitatively assessed. This explains why some of the keywords in our search string were not included in the final analysis (Fig 1).

#### 2.5.4 Subgroup analysis

As the 3-level models revealed significant heterogeneity, we conducted subgroup analyses to further investigate its source. In particular, we tested the influence of the technical moderators (type of ELS model and amount of potential bias) as well as of the brain regions within the brain areas previously described (Fig 3), in subsets of the dataset with sufficient observations. For information on the type of ELS model used, please see supplementary appendix S2-1.

#### 2.5.5 Sensitivity analysis and publication bias

According to the standards of meta-analyses, we should investigate the robustness of our effect sizes by performing analysis only on those studies that were blinded and randomized. Unfortunately, the amount of blinded and randomized studies was insufficient to proceed with this approach. As an alternative, we performed the analysis by including the amount of potential bias as a moderator. The results of this sensitivity analysis should be interpreted as the effects of ELS on biochemical markers of the dopaminergic system on studies of average risk of bias.

To detect publication bias, funnel plots’ asymmetry for each outcome variable was qualitatively evaluated. To the best of our knowledge, there are no available methods to quantify missing data (due for example to publication bias) in a multi-level setting(Assink and Wibbelink 2016). Nonetheless, we evaluated publication bias with Egger’s regression(Egger et al. 1997). However, these results should be interpreted with caution as they are not based on a 3-level model. Lastly, we excluded those studies responsible for funnel plot asymmetry and conducted sensitivity analysis on the remaining dataset in the attempt to evaluate the influence of publication bias in the meta-analysis.

Data are presented as *Hedge’s G* and 95% C.I. Data analysis was conducted with the computer program R (version 3.2.3)(Team 2015), with the aid of the following R packages: 1) *metafor*(Viechtbauer 2010) for conducting the analysis, 2) *ggplot2*(Wickham 2009) for graphics, and 3) shiny(Chang et al. 2017) to create the MaDEapp.

## 3 Results

### 3.1 STUDY SELECTION AND CHARACTERISTICS

#### 3.1.1 Study selection and data extraction

The process of study selection is illustrated in the flow chart (Fig 4). The search string identified a total of 979 unique research papers. Statistical measurements (e.g. mean, SD and N) for quantitative analysis were extracted from 90 peer-reviewed publications (that met our pre-specified inclusion criteria as described in the methods section. Three publications(Arborelius and Eklund 2007; Kippin et al. 2008; Kosten, Zhang, and Kehoe 2005) were excluded from the analysis as it was not possible to extract nor infer any statistical measurement. Similarly, information was lacking from 23 comparisons of three other publications (Kikusui, Faccidomo, and Miczek 2005; Ognibene et al. 2008; Kirsten et al. 2012).

The included studies dated between 1996 and 2016, used ~2600 animals yielding a total of 1009 comparisons from 152 experiments. The publications were analyzed in two separate datasets, respectively using prenatal (41 publications) and postnatal (49 publications) ELS models. For a summary of experimental characteristics across studies see supplementary Table S3-1.

Four observations of the prenatal dataset (striatal Th (Baier et al. 2014), striatal DA(Basta-Kaim et al. 2011), striatal HVA(Z. D.Ling et al. 2004), striatal DR2(Son et al. 2007)) and 6 observations of the postnatal dataset (striatal DR1(Y. Zhang et al. 2013), striatal DR2(Y. Zhang et al. 2015), DAT in the VTA area(Tien et al. 2013), striatal DOPAC(Ognibene et al. 2008), cortical DA(Matthews et al. 2001) and limbic HVA(Kubesova et al. 2015)) were excluded from the analysis as outliers.

#### 3.1.2 List of included publications

The publications included in the analysis were:(Adrover et al. 2007; Alonso, Navarro, and Rodriguez 1994; Baharnoori, Bhardwaj, and Srivastava 2013; Baier et al. 2014; Bakos et al. 2004; Basta-Kaim et al. 2011; Berger et al. 2002; Bingham et al. 2013; Bitanihirwe et al. 2010; Brenhouse, Lukkes, and Andersen 2013; Cabib, Puglisi-Allegra, and D’Amato 1993; Cai et al. 2013; Camp, Robinson, and Becker 1984; Carboni et al. 2010; Chocyk et al. 2011; Kwok Ho Christopher, Choy, De Visser, and Van Den Buuse 2009; K. H C Choy and van den Buuse 2008; Cory-Slechta et al. 2009; Cory-slechta et al. 2013; Dallé, Daniels, and Mabandla 2016; Delattre et al. 2016; L. W. Fan et al. 2011; L.-W. Fan et al. 2011; Gerardin et al. 2005; Gondré-Lewis et al. 2015; Gracia-Rubio et al. 2016; Granholm et al. 2011; Hall et al. 1999; Emily Hensleigh and Pritchard 2015; E. Hensleigh and Pritchard 2014; Hermel et al. 2001; Hill et al. 2014; J. W. Jahng et al. 2010; Jeong Won Jahng et al. 2012; Katunar et al. 2010; Kawakami et al. 2013; Kikusui, Faccidomo, and Miczek 2005; Kirsten et al. 2012; Kubesova et al. 2015; Lazzaretti et al. 2015; Lejeune et al. 2013; Li et al. 2014; Z. D. Ling et al. 2004; Z. Ling et al. 2004; Z. Ling et al. 2009; Liu et al. 2016; Lovic et al. 2013; Luchicchi et al. 2016; Madruga et al. 2006; Matthews et al. 2001; Meyer, Nyffeler, Schwendener, et al. 2008; Meyer, Nyffeler, Yee, et al. 2008; Novak et al. 2013; Ognibene et al. 2008; Oreland et al. 2011; Ozawa et al. 2006; Pallarés et al. 2013; Panagiotaropoulos et al. 2004; Pang et al. 2015; Papaioannou et al. 2002; Ploj, Roman, and Nylander 2003; Rentesi et al. 2013; Reynaert et al. 2016; Reznikov and Nosenko 1995; Romano-López et al. 2015; Eva Romero et al. 2007; E Romero et al. 2010; Rossi-George et al. 2011; Rots et al. 1996; Sasagawa et al. 2017; Silvagni et al. 2008; Silveira et al. 2010; Son et al. 2007; Tien et al. 2013; Vazquez et al. 2007; Vuillermot et al. 2012; Wang et al. 2009; Weston, Sobolewski, et al. 2014; Weston, Weston, et al. 2014; Winter et al. 2009; Womersley et al. 2011; Zager, Mennecier, and Palermo-Neto 2012; Zavitsanou et al. 2013; Y. Zhang et al. 2015; X. Y. Zhang, Kehoe, and Kosten 2006; Y. Zhang et al. 2013; X. Zhu et al. 2010; X. Zhu et al. 2011; Y. Zhu, Carvey, and Ling 2007; Zuckerman et al. 2003)

### 3.2 Meta-analysis: prenatal ELS

#### 3.2.1 Heterogeneity

Substantial heterogeneity was recorded in the prenatal dataset *(Q(378) = 954.969, p< .001)*, indicating that our search string identified a diverse range of experiments. In particular, 34.7% of variance could be attributed to within-sampling variance, 18.7% to within-experiment variance and 46.6% to between-experiment variance. These results suggested the use of moderators.

#### 3.2.2 Moderators

Potential moderators (supplementary Table S3-2) - such as brain area, sex, and species - were selected prior the beginning of the study based on hypotheses of the ELS literature.

As identified with univariate analysis of potential moderators (supplementary Table S3-3), outcome measure *(F(7,371)* = *3.956, p<.001)* and brain area investigated *(F(4,374)* = *6.144*, *p*<.*001)* were significant moderators in the prenatal dataset, explaining 4.3% and 6% of variance respectively. There was no detectable moderating effect of sex, species, age used, and method of assessment (RNA, protein or functional level).

#### 3.2.3 Model

The moderators that were univariately identified were included in the 3-level model to investigate the effects of ELS on markers of dopaminergic signaling. We hypothesized that the effects were dependent on the outcome analyzed and that they differed across brain areas.

Of the 20 interactions between outcome measure and brain area with enough data-points, only 2 reached statistical significance (Fig 5). For a summary of all interactions, see supplementary Table S3-4. In particular, in the striatal zone, Th was decreased *(Hedge’s G(se)* = *-1.164(.295)*, *p*<*.001*, supplementary Fig S3-1) while DOPAC was increased (*Hedge’s G(se)* = *.323(.136)*, *p* = *.018*, Fig 6A) following prenatal ELS.

**Fig 5.**
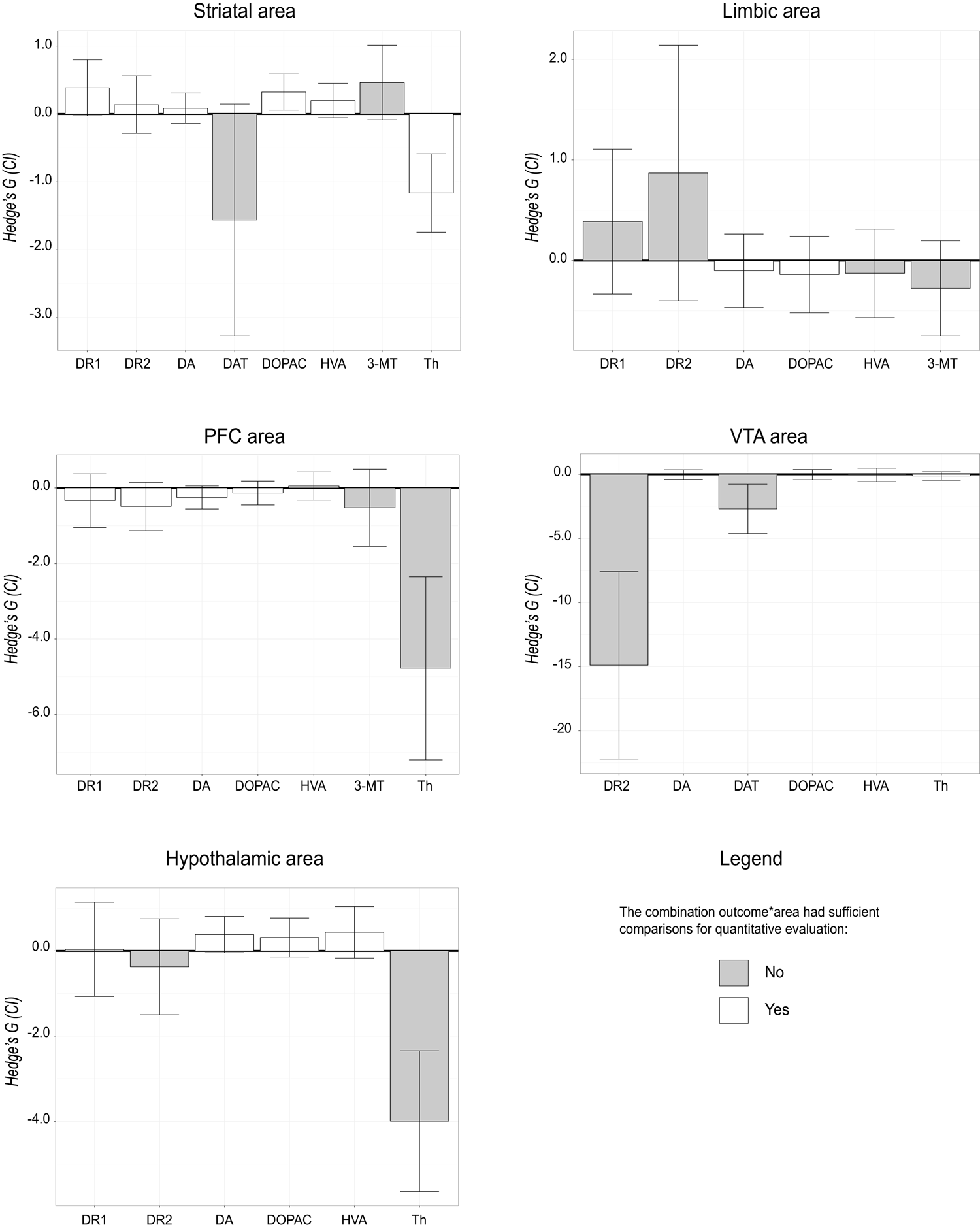
Summary of effects: prenatal dataset. Boxplot representing the summary of effect estimates for every combination between outcome variable (biochemical markers) and brain area. White bars = enough comparisons for meaningful quantification (rule of thumb: at least 4 comparisons from 3 papers), black bars = number of comparisons insufficient for analysis.

**Fig 6.**
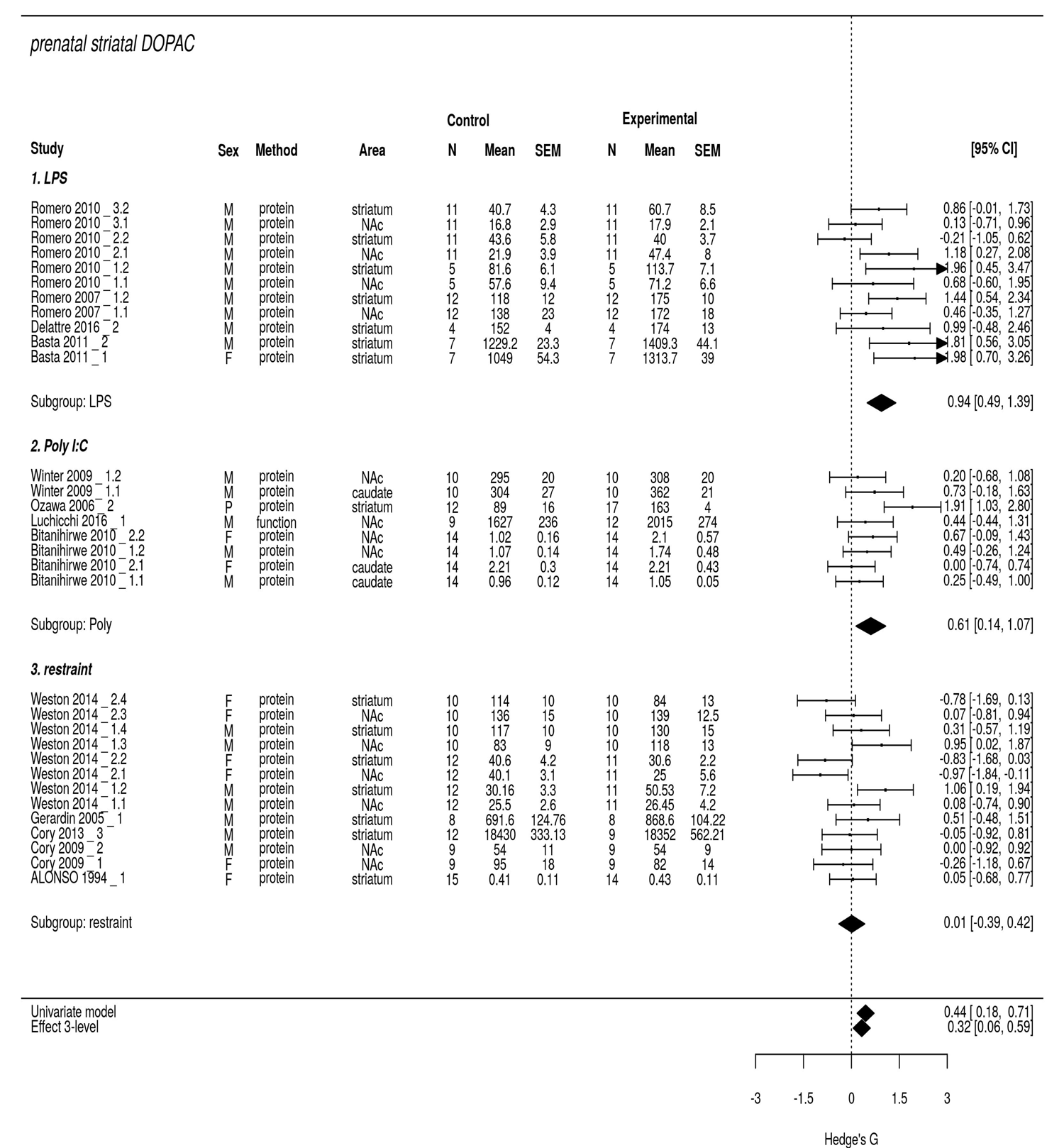
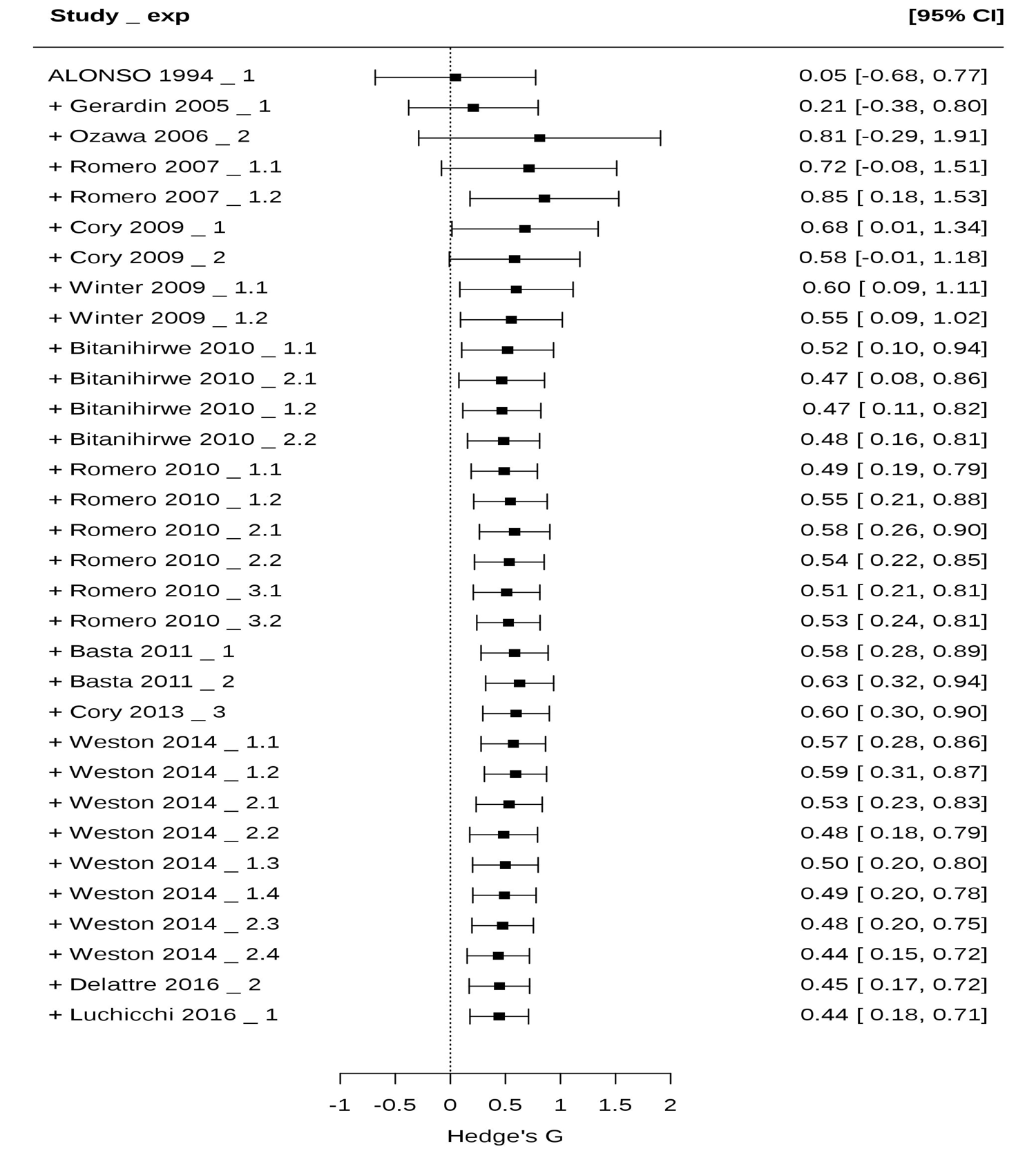
Striatal DOPAC in the prenatal dataset. (A) Forest plot. Results of univariate and 3-level meta-analysis are reported (bottom diamonds), as well as subgroup analysis on the ELS model used. (B) Cumulative forest plot. This plot displays the accumulation of evidence over time as the individual comparisons are added in chronological order. M = males, F = females, Method = method of assessment, hip = hippocampus, N = amount of animals, SEM = standard error of the mean, CI = confidence interval. Following the name of the study, _n represents the number of the experiment, .n = represents which comparison within that experiment.

Fig6B shows a cumulative forest plot of striatal DOPAC to exemplify that the chronological combining of the experiments shows consistency since 2010, and that subsequent experiments have not contributed to the direction nor the size of the effect. The cumulative forest plot does not correct for the multi-level structure of the model.

The interaction between outcome measure and brain area explained 22.2% of variance in the prenatal dataset. We identified 17 interactions with enough comparisons to further address heterogeneity *(Q(346)* = *742.97*, *p*<.*001)*.

#### 3.2.4 Subgroup analysis

Subgroup analysis was used to further investigate the unexplained heterogeneity deriving from the 3-level model. The type of ELS model and sub-brain areas were univariately evaluated as potential moderators for each interaction between outcome measure and brain area.

In the prenatal dataset, 15 interactions had enough observations to be considered for further subgroup analysis. Of these, 5 had a significant test of ELS model as moderator (supplementary Table S3-5), namely DOPAC and D1R in the striatal area, DA in the cortical area, and Th in the VTA area.

Subgroup analysis revealed that injection of LPS or PolyI:C has consistently different effects than maternal restraint. In particular, LPS and PolyI:C significantly increased DOPAC in the striatum *(LPS: Hedge’s G(se)* = *.941(.229)*, *p*<*.001*; *PolyI:C: Hedge’s G(se) = .608 (.238), p=.016)*, while restraint did not *(Hedge’s G(se)= .012(.207), p = .956, Fig 6A)*. Conversely, restraint decreased Th in the VTA area *(Hedge’s G(se) = -.85(.348), p=.03)* whilst LPS injection did not *(LPS: Hedge’s G(se) = .026(.188), p = .889*, supplementary Fig S3-2). Concerning cortical DA, the effects of PolyI:C and restraint had opposite directions but did not reach statistical significance, whilst the LPS model could not be quantitatively evaluated. Concerning D1R, no significant effects of the subgroup analyses were recorded.

Sub-brain area was a significant moderator only for DA in the striatal area (including nucleus accumbens, dorsal and central striatum and nucleus caudatus; supplementary Fig S3-3). In particular, DA was increased in the nucleus accumbens after ELS *(Hedge’s G(se) = .392 (.159), p = .01)* but it was unaffected in other parts of the striatum *(Hedge’s G(se) = -.122 (.146), p = .40)*. For a summary of all subgroup analyses, see supplementary Table S3-6.

### 3.3 Meta-analysis: postnatal ELS

#### 3.3.1 Heterogeneity

Our search identified a diverse range of experiments in the postnatal dataset, as shown by the substantial heterogeneity recorded *(Q(381) = 1061.278, p<.001)*. In particular, 33.1% to within-sampling, 46.9% to within-experiment, and 20.1% to between-experiment variance.

#### 3.3.2 Moderators

Univariate analysis of potential moderators (supplementary Table S3-7, *p*-value significance set at <.10) identified outcome measure *(F(8, 373) = 9.139, p < .001)* and brain area investigated *(F(4,377) = 2.035, p = .089)* as significant moderators, explaining 10.3% and 0.8% of variance respectively. Other moderators, such as species and sex of the animal, had no detectable moderating effect.

#### 3.3.3 Model

The moderators that were univariately identified were included in the 3-level model to investigate the effects of ELS on markers of dopaminergic signaling. We hypothesized that the effects were dependent on the outcome analyzed and that they differed across brain areas.

Of the 17 interactions with sufficient comparisons, 3 reached statistical significance (Fig 7, supplementary Table S3-8). In particular, in the striatal zone, DOPAC *(Hedge’s G(se) = .541(.207), p = .009*, Fig 8), HVA *(Hedge’s G(se) = .555(.22), p=.012*, supplementary Fig S3-4) and DA *(Hedge’s G(se) = .307(.147), p = .038*, supplementary Fig S3-5) were increased.

**Fig 7.**
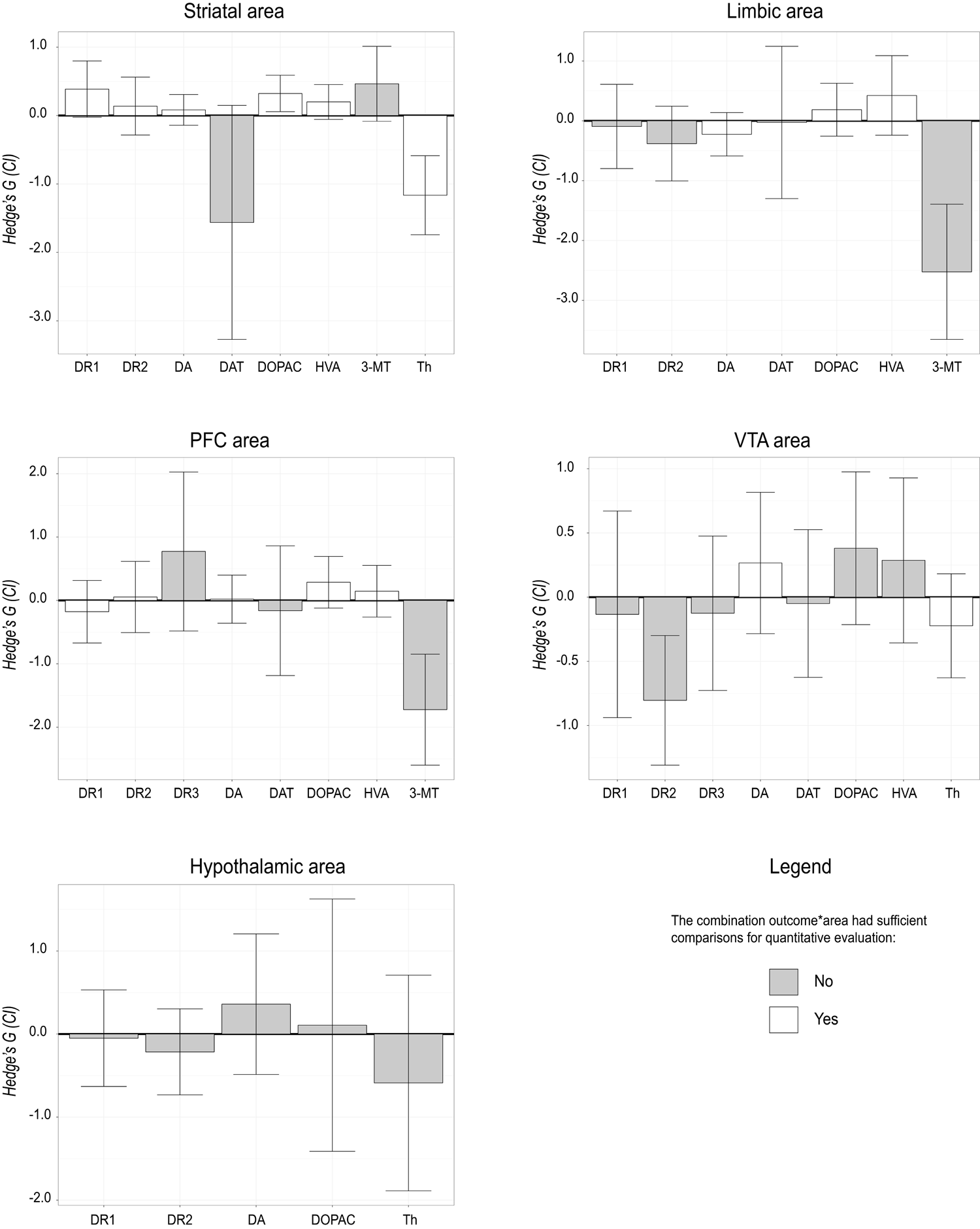
Summary of effects: postnatal dataset. Boxplot representing the summary of effect estimates for every combination between outcome variable (biochemical markers) and brain area. White bars = enough comparisons for meaningful quantification (rule of thumb: at least 4 comparisons from 3 papers), black bars = number of comparisons insufficient for analysis.

**Fig 8.**
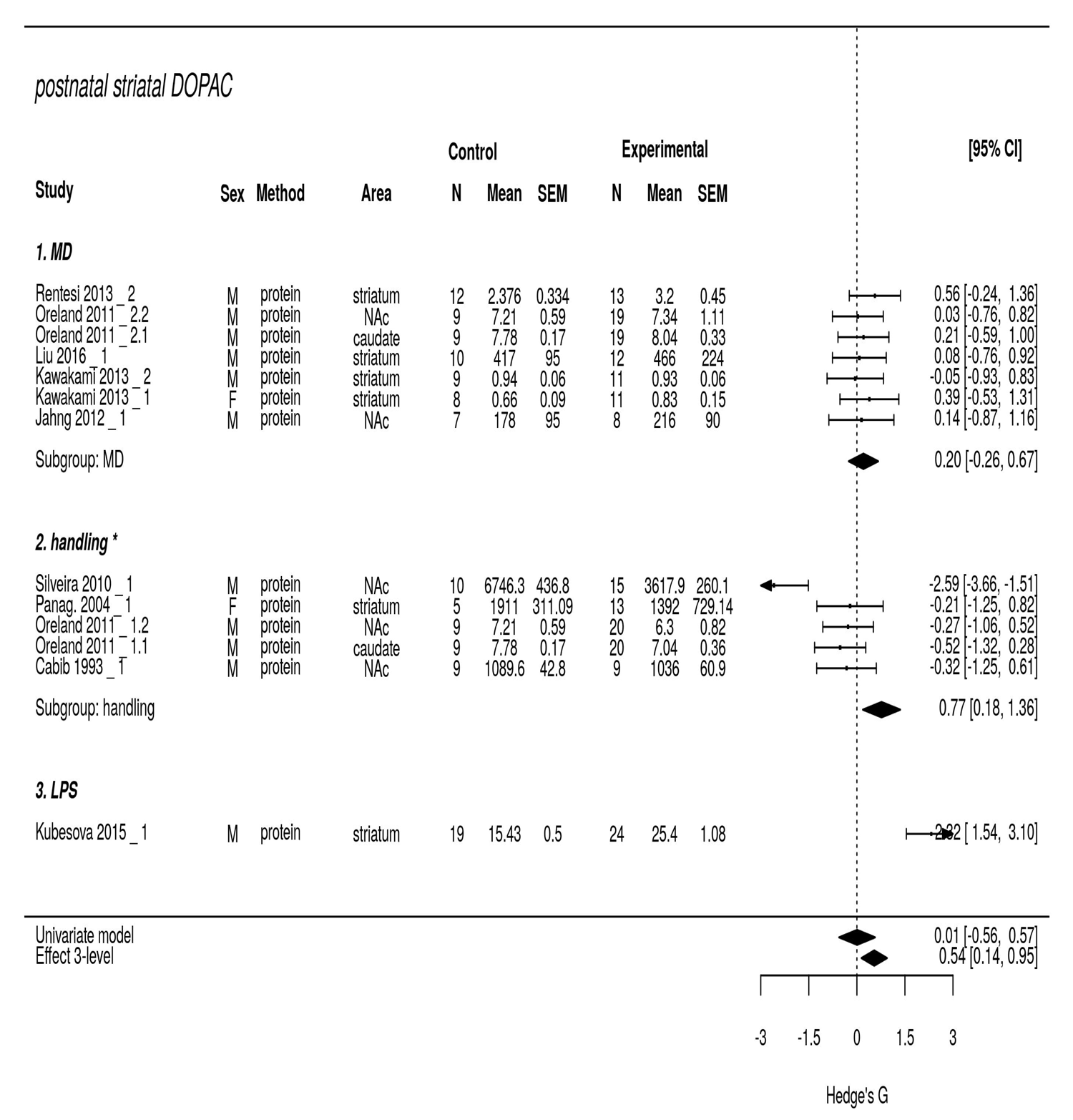
Striatal DOPAC in the postnatal dataset. Forest plot. Results of univariate as well as 3-level meta-analysis are reported (bottom diamonds), as well as subgroup analysis on the ELS model used. * = effect sizes of handling were multiplied by −1 to maintain consistency directionality of the other models. A decrease in the graph identifies an increase in DOPAC in ELS animals. M = males, F = females, Method = method of assessment, hip = hippocampus, N = amount of animals, SEM = standard error of the mean, CI = confidence interval, MD = mother separated from the pups, LPS = injection of LPS. Following the name of the study, _n represents the number of the experiment, .n = represents which comparison within that experiment.

The interaction between outcome measure and brain area explained 15.3% of variance in the postnatal dataset. We identified 16 interactions with enough comparisons to further address heterogeneity *(Q(345) = 898.4*, *p*<.001).

#### 3.3.4 Subgroup analysis

The moderator effects of the ELS model used and sub-brain areas were evaluated with a subgroup analysis.

In the postnatal dataset, 16 interactions had sufficient observations to be further analyzed. Of these, 7 revealed a significant impact of the ELS model used (supplementary Table S3-9): HVA, DOPAC and DA in the cortical area, HVA and DOPAC in the striatal area, DOPAC in the limbic area, and DA in the VTA area.

Subgroup analysis showed that the effects of ELS model as moderator in striatal DOPAC (Fig 8) and HVA (supplementary Fig S3-4) were mainly due to handling. In particular, handling decreased HVA *(Hedge’s G(se) = -.778(.295), p* = .03) as well as DOPAC *(Hedge’s G(se) = -.77(.301), p* = *.029*) in the striatum, whilst separation of the mother from the pups had no effect (*HVA: Hedge’s G(se) = .08(.227), p = .735; DOPAC: Hedge’s G(se) = -.205(.239), p = .411*).

Sub-brain area was not a potential moderator in any of the interactions evaluated.

For a summary of all subgroup analyses, see supplementary Table S3-10.

### 3.4 Sensitivity Analysis

Sensitivity analysis was conducted to test the robustness of our findings. We examined whether the quality of the studies included had an impact on the interpretation of our results.

#### 3.4.1 Quality of the studies: SYRCLE bias report

No publication reported information on all SYRCLE potential bias items. Overall, of the 90 publications, 37 (41%) reported randomization sequence of the animals in the experiments, 3 (3.3%) random housing allocation, 59 (65%) random group allocation, 49 (54.4%) random selection of the animals (Fig 9). In 11 (12%) publications the caretaker were reported blinded to the experimental condition, in 20 (22.2%) the experimenters blinded. Handling of incomplete data was reported in 42 publications (46.7%). 11 studies (12.2%) did not report sufficient information to evaluate the quality of the control group. Only 11 studies yielding a total of 117 comparisons reported being blinded and randomized.

**Fig 9.**
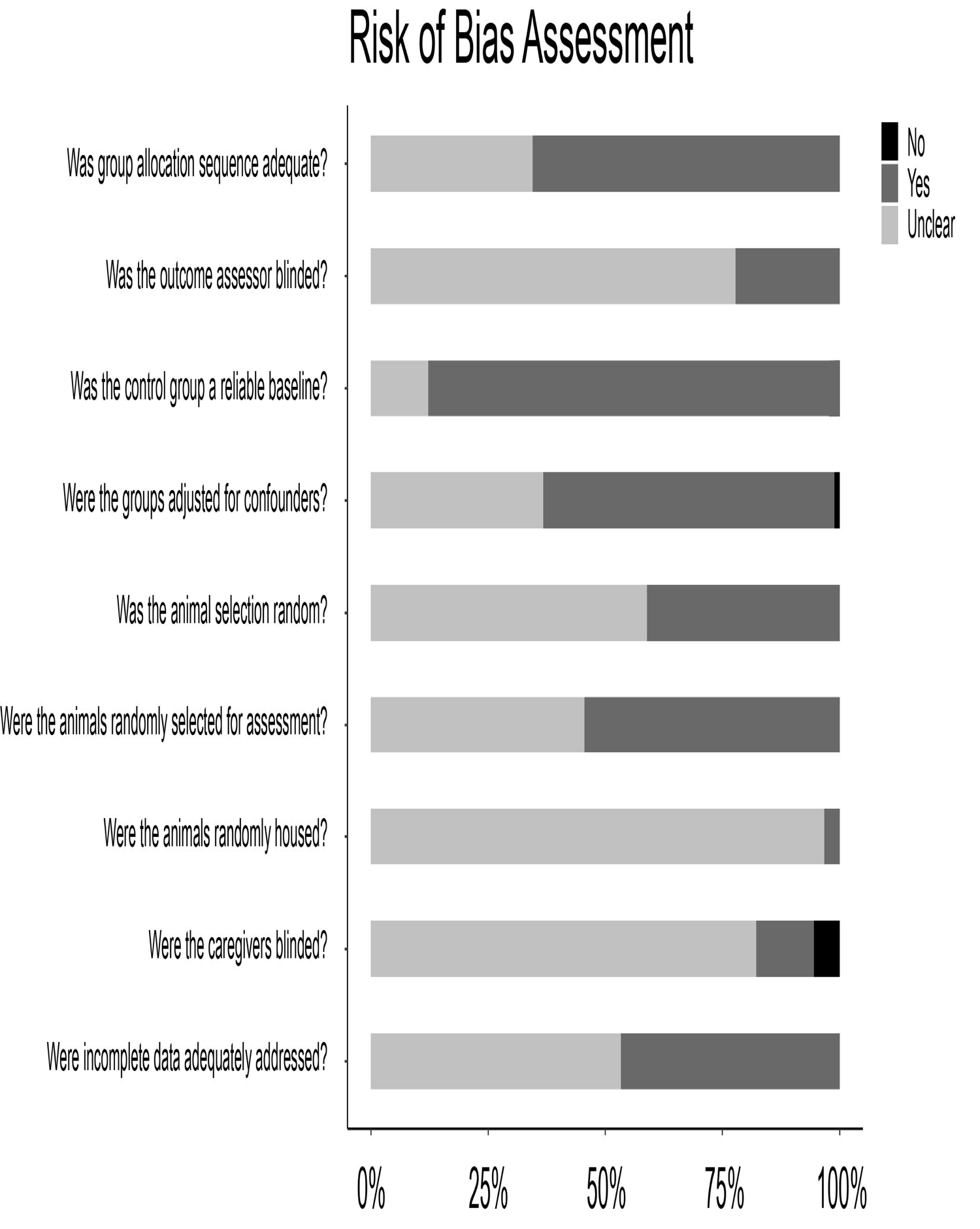
Risk of bias assessment. Each bar represents a different risk of bias item. Yes = measurements have been taken to avoid bias; no = no measurements were taken to avoid bias; unclear = not enough information provided in the paper to determine the risk of bias.

#### 3.4.2 Sensitivity analysis for potential bias

Since the amount of publications was insufficient to evaluate the robustness of our effects in a blinded and randomized dataset, we operationalized the amount of potential bias and performed the analysis again by including this factor as a moderator. Therefore, the results of this sensitivity analysis were interpreted as the effects of ELS on markers of DA signaling on studies of average bias.

The amount of potential bias was a significant moderator in the prenatal dataset (*F(1, 377)* = *3.536, p = 0.061*); yet, this did not affect the qualitative interpretation of the meta-analysis.

In the postnatal dataset, the test of moderators for amount of potential bias was not significant *(F(1*, *380)* = *0.500*, *p* = *0.480)*. The interpretation of the results did not change, with the exception of DA in the striatal area, of which the effect size was decreased and the effect at a trend level *(Hedge’s G(se)*=−.*289(.15)*, *p* = .*057*).

#### 3.4.3 Publication bias

Due to the lack of methods to quantitatively evaluate publication bias in a multi-level setting, we qualitatively estimated the risk for publication bias with funnel plots (Fig 10). Publication bias was more pronounced in the prenatal than the postnatal dataset. The same conclusion was reached when performing Egger’s regression (no multi-level regression models): there was evidence for publication bias in the prenatal (z = −5.014, *p* < .001) but not in the postnatal (z = −0.612, *p* = 0.54) datasets. The presence of publication bias in the prenatal dataset may indicate an overestimation of the reported effect sizes.

**Fig 10.**
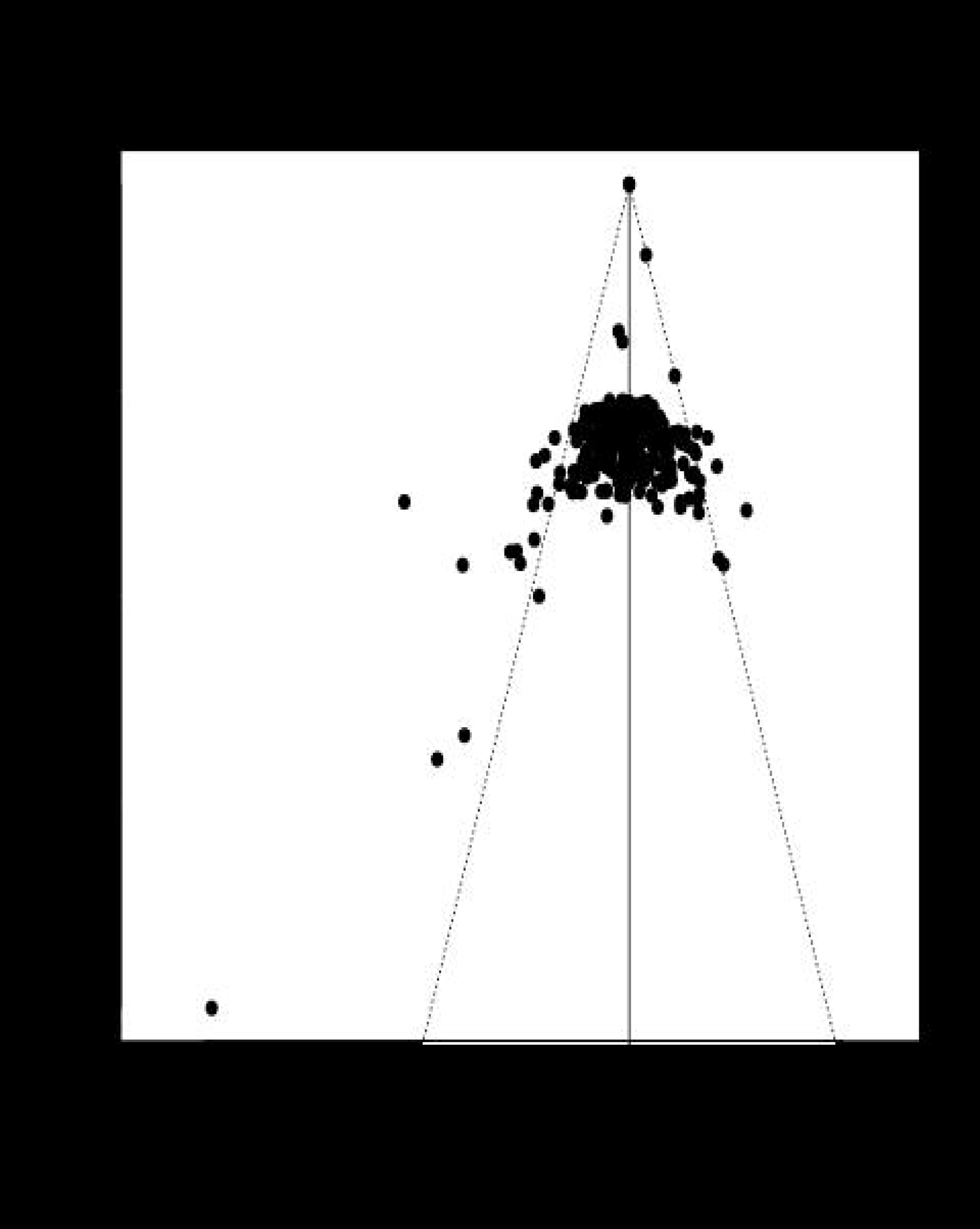
Funnel plots. Publication bias was evaluated by qualitatively assessing symmetry in funnel plot in the (A) prenatal and (B) postnatal datasets.

Furthermore, we conducted an analysis of influential cases by removing studies with large standard error as well as residual values. Since the results did not change qualitatively, publication bias was considered low-to-moderate.

### 3.5 MaDEapp

Finally, we created a MaDEapp ( https://vbonapersona.shinyapps.io/MaDEapp/), a web-based app with a user-friendly interface in which each researcher can perform his/her own meta-analysis on the topic of ELS and biochemical indicators of the dopaminergic signaling. The app offers the possibility to choose across a wide variety of options, such as outcome measures, brain areas, sex of the animals, type and timing of the ELS model. Based on the characteristics indicated, the app reports forest, funnel and cumulative plots. The forest plot includes a 3-level effect estimate (Hedge’s G and CI), which can be used for future power calculation.

For example, a researcher is interested in the effects of postnatal ELS on DR1 in the striatal area. In MaDEapp, the researcher selects the “postnatal” dataset, with “DR1” as outcome measure in the “striatal area” (Fig. 11). The resulting forest plot reports the estimated *Hedge’s G* (*CI*) = −.*5 [*-.*91*, −.*1]*. The estimated effect size is smaller than 0. From this exploration, the researcher hypothesizes that postnatal ELS decreases DR1 expression in the striatal area. The effect size −.5 may be an overestimation of the real size of the effect due to potential (publication) bias. The researcher would then use an effect size of −.45 for power calculation for his/her future experiments.

**Fig 11.**
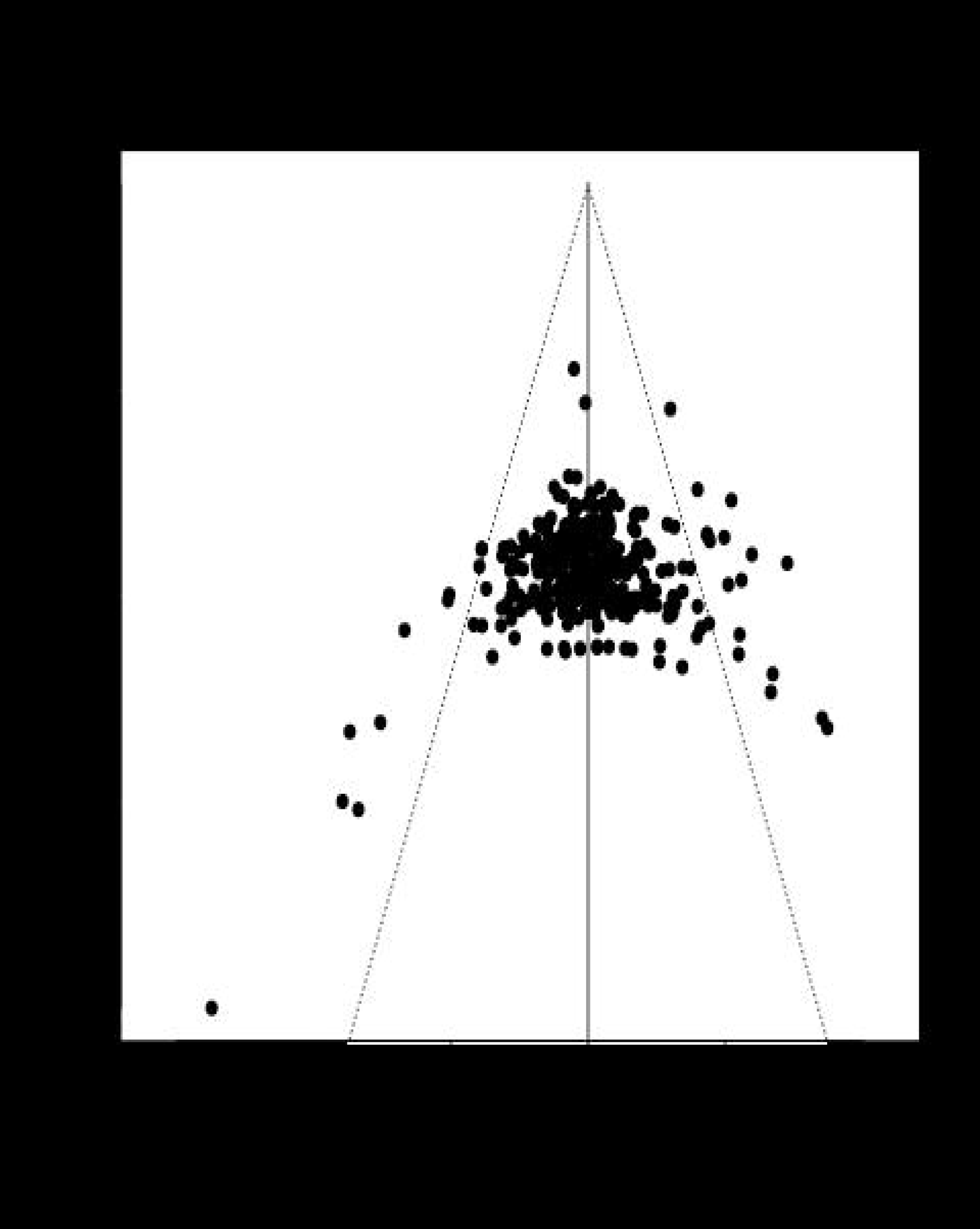
MaDEapp. Screen shot for visualization of MaDEapp, a web-based tool to perform ad hoc meta-analysis of the topic of ELS and biochemical indicators of the DA system. The app can be found at **website**.

## 4 Discussion

Schizophrenia and addiction are examples of psychiatric disorders reported to be linked to DA dysfunction. Childhood trauma is a well-documented risk factor(Teicher et al. 2016; Gatzke-Kopp 2011; Money and Stanwood 2013; Grace 2016). This clinical observation led to the hypothesis that the dopaminergic system mediates the risk of ELS. Although this link has been causally investigated in more than 90 rodent publications over 20 years, no consensus has yet been reached on the extent, directionality and specificity of this effect. Therefore, we performed a meta-analysis to question: Do rodent studies support long-lasting effects of ELS on biochemical indicators of the dopaminergic system? Overall, our results indicate that only a limited number of comparisons were significant suggesting that the effects of ELS on the dopaminergic system may not be apparent on a biochemical level.

### 4.1 Methodological considerations

Dopaminergic signaling involves multiple interdependent elements (e.g. precursors, metabolites, receptors), which altogether contribute to the system as a whole. Data on these elements are sometimes gathered from the same animals, and are therefore dependent on each other(Aarts et al. 2014). In a meta-analysis setting, dependency implies overlap of information, which ultimately leads to an erroneous interpretation of the results(Rosenthal 1991; Van den Noortgate et al. 2013). To deal with this obstacle, several strategies have been adopted: from selecting only one effect size per study to ignoring the problem altogether(Cheung 2014b). Although sophisticated methods such as multivariate and multilevel analysis exist, these have the strong limitation that the needed covariance between the dependent effects is rarely reported in publications(Hox 2010). The 3-level approach that we used overcomes both limitations: it corrects for dependency of observations, without the use of covariances(Assink and Wibbelink 2016). To the best of our knowledge, this approach has never been used before in rodent literature. Although this powerful and practical method was initially created for human studies(Cheung 2014b), its applicability in preclinical research is warranted due to the multiple-outcome nature of such studies. The method is already available in the R packages *metafor* and *metaSEM*.

Together with the application of a 3-level mixed effect meta-analysis to preclinical literature, we here promote the use of tools to facilitate data exploration and advocate open science. We created MaDEapp, a freely available user-friendly app that allows to run a tailor-made meta-analysis on ELS and the dopaminergic system, depending on the specific question one has. Each individual can select a set of characteristics (e.g. prenatal/postnatal models, sex, age). The app returns a forest plot in which the total univariate 3-level estimate is presented, as well as a funnel plot to evaluate publication bias. This can be used to generate hypotheses, evaluate estimated sample sizes for power analysis, and explore which outcomes/brain areas have received most attention and which did not. We believe this app is a useful tool to guide future research on the topic.

MaDEapp and the analysis here presented are complementary. Meta-analysis is a statistical test, and it is limited by the frequency (*power*) and quality (*potential bias*) of the published data. In our analysis there was no significant effect of postnatal ELS on DR1 in the striatum *(p* = .*053)*; however, when we analyzed the same outcome univariately with MaDEapp, the confidence interval of the estimate did not include 0 and could be interpreted as “significant”. This discrepancy may be due to a lack of power to confirm the effect in our analysis or due to an increased bias not corrected for in the more specific model used by the app. Therefore, non-significant results should be interpreted as lack of confirmation of an effect, not as evidence of no effect, since the meta-analysis could be underpowered to detect a specific marker in a specific brain area. Alongside, the use of the app should be intended as exploratory only and not as confirmatory.

### 4.2 Quality of the studies

Meta-analysis as a methodology is not simply the collection of statistical methods used to achieve integration of available evidence. Its power lies in the application of systematic scientific strategies to the literature review(Cornell and Mulrow 1999). In addition to summarizing effects’ estimates, it allows to evaluate the extent to which conclusions are at risk of bias.

In our analysis, surprisingly few studies (12%) reported being randomized as well as blinded. On the other hand, random allocation to group (41%) as well as blinded assessor (22.2%) was comparable(Antonic et al. 2013) or better(Egan, Sena, and Vesterinen 2011) than previous publications in neuroscience. Although it is likely that investigators did take measures to reduce bias, lack of their reporting induced an unclear risk(Kilkenny et al. 2013; Landis et al. 2012) and hindered estimation of the value of the publications(Kilkenny et al. 2013). The importance of quality of reporting has been an emerging issue in preclinical research(Kilkenny et al. 2013; Landis et al. 2012). Despite the increased awareness, the quality of reporting of the publications included in this meta-analysis has not improved since 2005 (supplementary Fig S4-1). Such evidence should encourage preclinical researchers as well as reviewers to adhere to reporting guidelines such as ARRIVE(Kilkenny et al. 2013).

Although imprecise reporting does not necessarily imply poor study quality, underpowered experiments seriously hamper research interpretation(Button et al. 2013). From the reported amount of animals included per experiment, we back-calculated the power at the beginning of the study, assuming at least one true positive effect per publication. We performed this analysis considering small (*Hedge’s* G = .5), medium (*Hedge’s* G = .8) or large (*Hedge’s* G = 1) effect sizes (supplementary Fig S4-2). When considering a large effect size, 391 comparisons (38.7%) had power below chance level, and only 63 (6.2%) had power >.8, a cut-off value(Cohen 1992) generally aimed at in preclinical research. Although 43 papers (47%) had at least one comparison with power >.5, only 5 papers (5.5%) had at least one comparison with power >.8. This means that the vast majority of the experiments was not sufficiently powered to reliably detect an effect – an (already dramatic) best-case scenario given that the percentages were calculated assuming that the studies compared only two groups (t-tests) as well as a large and truly existing effect. Future preclinical studies should be grounded in power calculations based on realistically estimated effects. Although for each single study the amount of animals will be larger, overall higher power will lead to more reliable, reproducible and therefore higher quality research.

### 4.3 ELS causes limited alterations on biochemical markers of the DA system

In our analysis, we evaluated the dopaminergic system by including numerous biochemical markers across brain areas as well as potential moderators. These gave rise to a myriad of viable comparisons. Despite the extent, only a handful of significant effects were identified, thereby suggesting that biochemical indicators of the DA system well adapt to ELS interventions. Clearly, we cannot exclude the possibility that other indicators of the DA system (e.g. electrophysiological parameters or behavioral tests that critically depend on DA function) would have yielded clearer results. This awaits future investigation.

Prenatal and postnatal ELS were treated separately because the prenatal environment differs substantially from that postnatally. Nonetheless, both datasets shared consistent findings. Specifically, the striatal area was the most vulnerable: following prenatal ELS, Th was decreased and DOPAC increased; while postnatal ELS caused an increased in DOPAC as well as HVA. These changes were stable and reliable: the analysis used is adequately conservative and robust, and the effects survived sensitivity analysis and publication bias corrections. The stability of the effects is also qualitatively substantiated by the cumulative plots, which operationalize how subsequent experiments update our knowledge of the previously estimated effect size. Our results display that these were durable over time, and that replication after the initial 3-5 studies might not be very informative on these variables (except as a positive control in a study investigating another variable), as additional experiments did not alter the estimated effect. All in all, the sparse effects here reported are reliable and of medium size, suggesting that the system is damaged, which may in turn contribute to the vulnerability of ELS-dependent disorders.

The results can be interpreted as either hyper- or hypo-activation. For instance, the increase in metabolites can indicate an increase in the available amount of the substrate DA (hyperactivation) as well as an increased conversion rate causing less DA (hypoactivation). Similarly, the decrease in Th (precursor conversion enzyme) is not necessarily indicative of a decrease in DA function. Although our analysis suggests that postnatal ELS increases DA levels in the striatum, the effect is small in size and less robust that the other effects mentioned above. The mismatch between DA precursor and metabolism may suggest changes in the intermediate stage of DA conversion. For example, the DA converting enzyme COMT has been repeatedly shown to interact with ELS for the later development of psychiatric disorders(Lovallo et al. 2017; Klaus et al. 2017; Sheikh et al. 2017). On the other hand, L-DOPA – product of Th and precursor of DA – has been suggested to act as a novel transmitter itself or may have neurotropic functions, and thereby be transiently involved in perinatal developmental processes(Braun et al. 1999). Since the interaction between ELS and L-DOPA has not been further investigated, the link remains circumstantial.

We defined a priori several factors established in preclinical literature to be potential moderators of ELS effects. To our surprise, species, sex and method of assessment were not significant moderators. Although males, mice and protein as method of assessment were the most described conditions, plenty of observations were present for all groups. Nonetheless, the lack of evidence for a moderator effect should not be interpreted as evidence for absence: mice/rats should be chosen according to standard practice, both sexes should always be considered(Bale and Epperson 2015), and there is substantial evidence that a decrease in RNA level does not automatically result in a decrease in protein and therefore in function, as e.g. shown in a systems approach(Williams et al. 2016).

Lastly, the unexplained variance may not only indicate methodological differences, but also underlie additional biological moderators, such as sub-brain areas or differences across hemispheres (lateralization).

### 4.4 Translational potential?

The translational applicability of preclinical studies depends on the understanding of psychopathological clinical and intermediate phenotypes. For example, ELS is a main risk factor for schizophrenia as well as substance abuse disorder. However, these diseases have opposite intermediate phenotypes: while schizophrenia is supposed to be characterized by hyperreactivity of the DA system although presumably to its afferent control(Grace 2016), substance abuse may be linked to DA hypo(re)activation(Wise and Koob 2014). To what extent do ELS studies in rodents accurately model these two conditions?

Three factors currently limit answering this question. Firstly, ELS in humans is a complex concept, generally involving low socio-economic status, physical and/or psychological abuse, poor living conditions and high caloric food (Teicher et al. 2016). Conversely, animal models are extremely controlled and standardized pre- and postnatally. Although this facilitates the definition of “traumatic early life” as well as the deriving caused effects, one can question its ecological validity. Secondly, the dual hypo- / hyper-interpretation of the ELS-induced phenotype in rodents prevents a whole-system level comparison, and restricts it to a micro level, focusing on a particular compound in a particular brain area. Thirdly, the DA-dependent changes in schizophrenia and addiction are most likely far more complex than the ones observed following ELS in rodents. For example, our analysis failed to confirm any effect of ELS on DA receptors. This was surprising, as changes in the availability of DA receptors is a consistent characteristic across different types of addictions(Nutt et al. 2015) as well as in schizophrenia(Sanyal and Van Tol 1997). Although the discrepancy could partly be due to a power problem of the meta-analysis, these limitations challenge the reliability of ELS models for translational purposes, at least with regard to these specific aspects of the abovementioned human disorders. It cannot be excluded that more relevant models may become apparent in light of different ELS theories(Daskalakis et al. 2013).

Lastly, although our study supports that ELS causes some changes in the DA system, these associations remain at a correlational level in humans and should be interpreted as such.

### 4.5 Limitations of the study

Despite our efforts to be as comprehensive as possible in the description of the effects of ELS on the DA system, we encountered several important limitations. Firstly, we investigated the DA system by evaluating the effects of ELS on biochemical markers. Although this provides a thorough conceptualization of the system, it does not supply a comprehensive functional evaluation. For example, the approach here reported is unable to operationalize DA innervations, projections and tone. ELS has been reported to alter DR3-signalling and neuronal activity in the lateral septum(Shin et al. 2018). Chronic stress in adulthood has been reported to change DA neurons’ activity in a stressor-dependent manner(Valenti, Gill, and Grace 2012). These reports suggest that spontaneous activity, bursting and timing of dopaminergic firing may be susceptible to ELS action, yet they are not apparent from assessment of ligands, receptors and metabolites. Despite the high relevance of such measurements, these were excluded from the analysis as the publications on the topic are scarce and their integration not straightforward in a meta-analytic setting.

Secondly, the classification of “timing” of ELS to either prenatal or postnatal may be too reductionist, since neuronal circuits are shaped by experiences during critical periods of development of variable length (from days to years depending on the species) (Hensch 2005). The interested researcher can further explore this avenue by combining our dataset with RNA expression of Th or DA receptors found in the Allen Developing Mouse Brain Atlas(Thompson et al. 2014). Unfortunately, the literature so far published is insufficient to investigate how stress during specific postnatal days in which a certain RNA X is highly expressed uniquely alters its functioning later in life.

Thirdly, due to insufficient data-points per outcome per brain area in several cases, a meaningful quantitative estimation was not possible for all combinations of outcome*area. All currently available measurements are reported as supplementary material and can be further investigated via MaDEapp.

Fourthly, we included data only from published studies. Especially in the prenatal dataset, there is evidence of low-to-medium publication bias as qualitatively estimated with a funnel plot and sensitivity analysis, which may result in an overestimation of the effect sizes. Despite the robustness of our methodology, this limitation should be considered in future power calculations.

Lastly, we limited our analysis to baseline (i.e. unchallenged) conditions. Future studies should focus on conditions where the DA system is challenged, as ELS manipulations can interact with later life challenges to result in a pathological phenotype(Daskalakis et al. 2013).

To conclude, ELS induces a few yet robust effects on biochemical indicators of the DA system, with – based on the currently available studies – the striatum being the brain area most affected. Although the changes observed can be interpreted as both hypo- and hyper-activation of the DA system, the effects were consistent across prenatal and postnatal ELS models, sex, species and method of assessment.

## 5 ACKNOWLEDGMENTS

This work was supported by the Consortium on Individual Development (CID), which is funded through the Gravitation program of the Dutch Ministry of Education, Culture, and Science and Netherlands Organization for Scientific Research (NWO grant number 024.001.003). R.A.S. was supported Netherlands Organization for Scientific Research (NWO Veni grant 863.13.02).

We thank Prof. Joop Hox for his input on multilevel methods, and Erik-Jan van Kesteren for his programming expertise. We also gratefully acknowledge the in depth discussions and feedback on the methods by Prof. Herbert Hoijtink and Dr Remmelt Schür.

